# Calcium transients regulate epithelial integration of multiciliated cells during Xenopus skin development

**DOI:** 10.1101/2024.11.01.621480

**Authors:** Neophytos Christodoulou, Paris A. Skourides

## Abstract

Integration basally located progenitors into an existing epithelium is crucial for tissue morphogenesis during embryogenesis and tissue homeostasis of epithelial organs. In this study, using Xenopus as a model system, we explore the role of intracellular calcium in multiciliated cell (MCC) progenitors’ epithelial integration during mucociliary skin epithelium development. Our findings reveal that calcium transients precede MCC apical emergence and are essential for their successful insertion into the overlying skin epithelium. Furthermore, we demonstrate that phospholipase C (PLC) activity is required for the generation of calcium transients, which in turn regulate MCC epithelial integration via the calcium-binding protein calmodulin. Last, we demonstrate that the PLC/Ca²⁺/calmodulin axis is necessary for proper formation of the apical actin network, by influencing its stability. This study advances our understanding of the molecular mechanisms governing epithelial integration of basal progenitors and highlights the importance of calcium signalling in coordinating cytoskeletal dynamics during epithelial morphogenesis.

## Introduction

Regular addition of new cells and removal of old ones is necessary for epithelial morphogenesis during embryogenesis and the homeostasis of epithelial organs^1^. New epithelial cells can arise from cell divisions within the epithelial sheet^2,3^ or from a pool of basally localized progenitors^4–6^. These progenitor cells give rise to new epithelial cells that join the existing epithelial layer through a series of coordinated actions. Initially, the basal progenitors acquire apicobasal polarity^7^, intercalate through the basolateral region of epithelial cells^5,6^, anchor at the apical junctions of the epithelium and remodel those junctions to join the epithelium^8^ and expand their surface area^9,10^, to a size accommodating the cell function^11^. Insertion of basally localized progenitors into an overlying epithelium has been shown to contribute to embryogenesis ^5,12–14^ and tissue homeostasis^4,15–19^.

The study of the molecular and cellular mechanisms regulating epithelial integration of basally situated cells is challenging, due to the complexity of the process and the inaccessibility of tissues presenting this behaviour. Multiciliated cell (MCC) epithelial integration during mucociliary skin epithelium development of Xenopus embryos has proven to be an invaluable model and has contributed to our understanding of epithelial integration^5,6^. Research on Xenopus mucociliary epithelium development has shown that MCC epithelial insertion is controlled by progenitor cell differentiation^20^, apicobasal polarity establishment ^7,21^, probing of the mechanical environment^8^, centriole amplification^22^, microtubule post-translational modifications^23^, actin network polymerization^9^ and stabilization^24^.

The apical actin network of MCCs has a central role in MCC epithelial insertion. Specifically, during MCC epithelial integration an apical actin network forms a lattice that generates the cell-autonomous forces required for epithelial insertion and apical surface expansion^9,10^. The apical surface of MCCs hosts hundreds of basal bodies and cilia^25^. As a result, tight regulation of apical actin network architecture and dynamics is essential for the apical emergence of MCCs and their subsequent function in the tissue^11^. Nevertheless, our knowledge regarding the molecular regulation of actin network assembly and stabilization during integration of new cells into an existing epithelium is still limited.

Calcium is a highly versatile intracellular messenger regulating a wide array of cellular processes by modulating the activity of calcium-binding proteins. For example, calcium affects actomyosin contractility through the regulation of myosin activity, contributing to processes such as cell migration^26^ and cell shape changes during embryogenesis^27–29^. In addition to its involvement in actomyosin contractility, it has been demonstrated that calcium regulates actin cytoskeleton architecture and dynamics. Specifically, calcium has been implicated in the regulation of actin filament formation, F-actin bundling and severing^30^. The bundling and severing of F-actin is controlled via direct interaction of actin-binding proteins with calcium^30^. In contrast, F- actin filament formation is regulated by the calcium-sensing protein calmodulin^30^. Upon calcium influx calmodulin changes conformation and interacts with several cytoskeletal effectors controlling downstream actin filament formation^31–33^. The role of the Ca^2+/^calmodulin signalling axis during basal progenitor insertion into a polarized epithelium remains to be elucidated.

Here we study the role of intracellular calcium in MCC epithelial integration in the Xenopus mucociliary skin epithelium. Our work reveals that intracellular calcium transients precede MCC apical emergence. Subsequently, we show that calcium transients are indispensable for MCC insertion in the skin epithelium and subsequent expansion of their surface area in a tissue-autonomous and cell-autonomous manner. Furthermore, we show that PLC activity is necessary for calcium transient generation and MCC epithelial integration. Additionally, we show that Calmodulin is an effector of intracellular calcium during MCC apical emergence, and its activity is crucial in this process. Last, we show that the PLC/Ca^2+^/Calmodulin signalling axis regulates apical actin enrichment during MCC apical emergence by influencing the stability of the actin network.

## Results

### Calcium transients precede MCC epithelial integration

MCC epithelial integration requires the generation of 2D pushing forces from an apical actin network^9^. Calcium actin interplay is crucial for various cellular processes during development and disease^27,28,34–36^. Thus, to examine a possible involvement of intracellular calcium during MCC epithelial integration we decided to image Ca^2+^ levels using GECO-RED, a genetically encoded Ca^2+^ indicator ^27,37^. The two ventral blastomeres of four-cell-stage Xenopus embryos were injected with mRNAs encoding membrane-GFP and GECO-RED to target the developing skin epithelium (**Figure 1A**). Subsequently, embryos were allowed to develop to stage 18 and imaged during MCC epithelial integration (**Figure 1A**). Live imaging revealed that intercalating cells frequently displayed transient Ca^2+^ level increases (**Figure 1B-C, Movie 1**). These Ca^2+^ transients were cell-autonomous and asynchronous with a duration of 20 seconds (**Figure 1D, Figure S1A**). To examine if the cells displaying calcium transients are intercalating MCCs, we cloned the calcium sensor GECO-RED under the control of the MCC-specific a-tubulin promoter^5^. Injection of linearized DNA plasmid encoding atub:GECORED revealed that the cells displaying calcium transients are indeed intercalating MCCs (**Figure 1E, Movie 2**). Subsequently, we went on to examine the temporal correlation between calcium transients and apical cell surface expansion in MCCs. Thus, we quantified calcium levels and apical cell surface area over time in intercalating MCC. This revealed that during MCC epithelial integration calcium transients precede MCC apical surface area expansion (**Figure 1F-F’’, Figure S1B, Movie 3**). In addition, quantification of MCC apical cell surface area increase over time in the absence or the presence of a calcium transient revealed a significant correlation between calcium transients and MCCs apical cell surface expansion (**Figure 1G)**. In support of the latter, live imaging of intercalating MCCs before their integration in the overlying skin epithelium, revealed that calcium transients appear only in MCC inserting into the skin epithelium (**Figure 1H**). Collectively these results show that Ca^2+^ transients precede MCC apical emergence (**Figure 1I**), suggesting that Ca^2+^ transients contribute to MCC epithelial integration and apical surface expansion.

**Figure 1.**
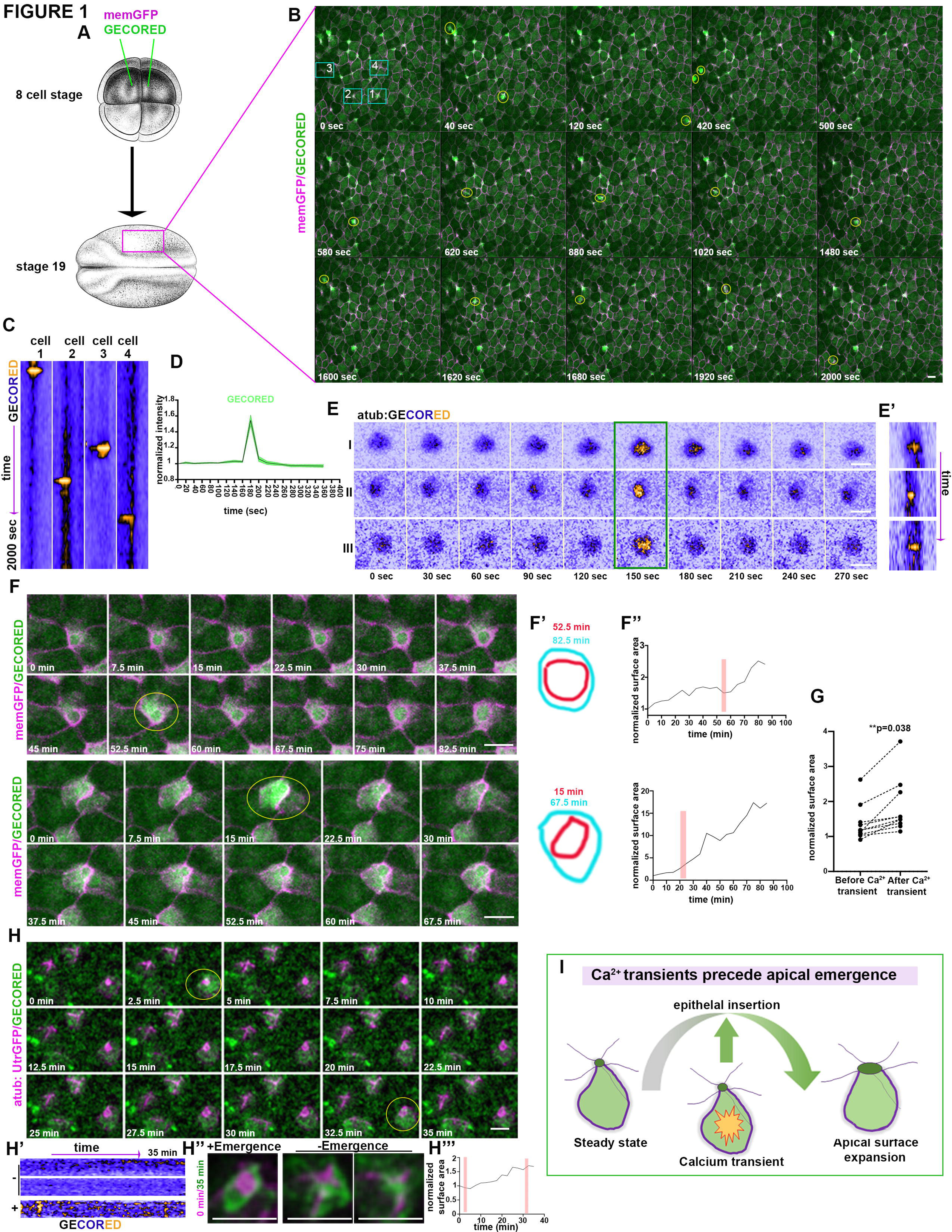
Calcium transients precede MCC epithelial insertion. A) Diagram of the experimental procedure followed for the generation to time-lapse recording. B) Stills from a time-lapse recording of an embryos expressing mem-GFP and the calcium sensor GEC-RED. Transient increases in calcium levels are indicated by yellow circles. C) Kymographs showing calcium levels over time in cells 1-4 marked by rectangles in B. D) Normalized GECO-RED intensity over time for distinct calcium transients. n=15 cells. The graph follows calcium levels 180 sec before and after each transient. E) Stills from representative MCC expressing atub: GECO-RED, 120 sec before and after the appearance of a calcium transient. E’) Kymographs of MCCs shown in E. F) Stills from time-lapse recording showing 2 representative intercalating MCC. Upon the appearance of calcium transients, cells expand their surface area. F’) MCC apical surface area during and after a calcium transient. F’’) Quantification of the surface area over time for MCC shown in F. Red rectangle represents the calcium transient. G) Quantification of MCC surface area increases in a 30min time window before and after the appearance of calcium transient. n=10 cells, two-sided, unpaired Student’s *t*-test. H) Stills from a time lapse recording following 3 MCC. Calcium transients (yellow circle) appear only in the apically emerging MCC. H’) Kymographs for atub:GECO-RED in the three MCCs shown in H. H’’) MIP images of the first and last time points for MCCs shown in in H showing successful epithelial integration of the MCC displaying calcium transients. G) Schematic for the temporal relationship between calcium transients and MCC apical emergence.

### Calcium transients are necessary for MCCs epithelial integration

Intracellular calcium transients display strong spatiotemporal correlation with MCC apical emergence. Thus, we decided to functionally assess the role of intracellular Ca^2+^ transients in MCC epithelial integration. To achieve this, we employed 2- aminoethoxydiphenyl borate (2APB), which has been demonstrated to impede IP3R mediated calcium release and efficiently blocks calcium transients in Xenopus embryos without affecting basal Ca^2+^ levels^28^. MCC start to integrate the skin epithelium at stage 18 and all MCCs are inserted into the skin epithelium by stage 24^5,23^. Thus, we treated embryos with 2APB from stage 15, and we imaged calcium levels in intercalating MCCs, revealing that 2APB efficiently blocks calcium transients in MCCs (**Figure S1C-E**). Subsequently, we treated embryos with 2APB from stage 15, before initiation of MCC epithelial integration and allowed the embryos to progress to stage 24, when MCC epithelial integration is completed. Subsequently, we assessed the effect of 2APB on MCC epithelial integration. In control embryos, MCCs were successfully inserted into the superficial skin epithelium (**Figure 2A-B**). In contrast, in 2APB-treated embryos MCCs failed to integrate into the superficial skin epithelium (**Figure 2B**). 2APB targets ER-mediated calcium release. ER-mediated calcium release has been implicated in TJ biogenesis^38,39^. Therefore, we went on to examine if the observed phenotype stems from disruption of the skin epithelium due to 2APB treatment. For this, we assessed the localization of adherens (β-catenin, E-cadherin) and tight junction markers (ZO-1). This revealed that adherent and tight junctions in 2APB treated embryos were unaffected (**Figure S2A-F**), indicating that observed phenotype stems from a direct disruption of intracellular calcium signalling rather than a generalized impairment of epithelial integrity.

**Figure 2.**
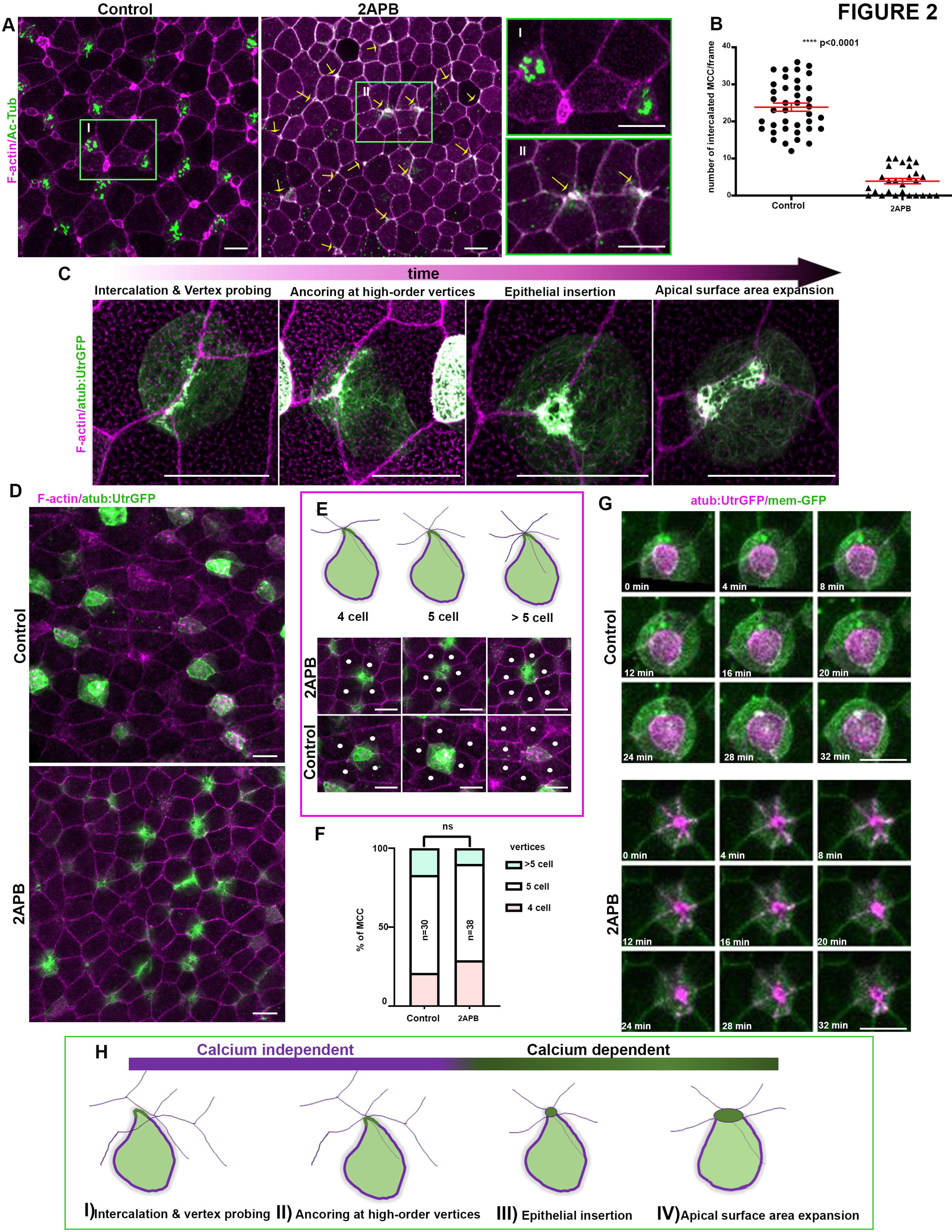
Calcium transients are indispensable for MCC apical emergence. A) Representative image of control and 2APB treated skin epithelium of stage 24 embryo. MCCs have successfully inserted into the skin epithelium of control embryos. MCCs epithelial integration is defective in 2APB treated embryos, with MCCs failing to acquire an apical surface area (arrows). B) Quantification of MCC epithelial insertion. n=39 positions from 24 control embryos and 27 positions from 15 2APB treated embryos. ****p<0.0001, two-sided, unpaired Student’s *t*-test C) Main steps of MCC epithelial insertion. D) Representative images of control and 2APB treated stage 24 embryos expressing atub:UtrGFP. E) Representative images of MCCs in control and 2APB treated embryos showing anchoring at high-order vertices. F) Quantification of the vertex type occupied by MCCs in control and 2APB treated embryos. χ^2^ test shows no significantly statistical differences (ns). G) Stills from time lapse recordings depicting MCCs upon their binding to high order vertices. MCCs in control embryo successfully inserts into the overlying skin epithelium expanding its surface area. The MCC in the 2APB treated embryo displays defective epithelial insertion after its binding to a high- order vertex. H) Schematic regarding the role of intracellular calcium in the distinct steps of MCCs epithelial insertion.

MCC epithelial integration is accomplished via distinct independent steps (**Figure 2C**). Upon their specification, MCCs initially intercalate from the deep epidermal layer towards the superficial layer and are positioned between the lateral side of superficial cells (**Figure 2C**). Subsequently, MCCs probe the mechanical properties of the overlying epithelium junctions and actively remodel the overlying junctions to generate high-order vertices of increased tension and anchor at these vertices (**Figure 2C**). Next, MCCs integrate into the skin epithelium and to complete the process, MCCs expand their surface area to a size supporting optimal cell function (**Figure 2C**). To identify which steps of MCC epithelial integration require intracellular calcium transients, we injected embryos with atub:UtrGFP to visualize MCCs, allowed embryos to develop to stage 15 and treated them with 2APB up to stage 24. This revealed that inhibition of calcium transients affects specific stages of MCC epithelial integration. Specifically, anchoring at high-order vertices was unaffected (**Figure 2 D-F**). In contrast epithelial integration and apical surface area expansion were defective as revealed live imaging of MCCs in control and 2APB-treated embryos (**Figure 2G, Movie 4**). In both conditions, MCCs anchor at an overlying high-order vertex. While MCC in the control embryo successfully insert into the overlying skin epithelium and expands its apical cell surface, the MCC in 2APB treated embryo fails to insert into the epithelium (**Figure 2G, Movie 4**). The above data reveal that Ca^2+^ transients contribute to specific steps during MCC epithelial integration (**Figure 2H**). Specifically, Ca^2+^ transients are dispensable for radial intercalation between the lateral region of superficial epithelial cells as well as the mechanical probing of apical junctions and the remodelling of these junctions. On the other hand, our data indicate that intracellular calcium transients are indispensable for MCC epithelial integration and apical cell surface expansion (**Figure 2H**).

### Calcium transients are involved in MCC epithelial insertion in a cell autonomous manner

During MCC epithelial integration, the skin epithelium receives forces from the underlying extending mesoderm. Mechanical tissue coupling has been shown to affect tissue morphogenesis in different models^40–44^. Specifically, stretching forces generated by mesoderm morphogenesis directly impact the planar cell polarization of MCCs^45,46^ as well as the size of MCCs^11^. Calcium transients have been shown to drive mesoderm convergent extension^47^. To exclude the possibility that mesoderm defects affect mucociliary epithelium development in embryos treated with 2APB, we decided to examine the effect of 2APB treatment in ectoderm (animal cap) explants^48^. These explants differentiate autonomously into a functional mucociliary epithelium and effectively replicate the initial stages of skin epithelium morphogenesis^49^. Animal cap explants were dissected from stage 9 embryos and cultured until the sibling embryos reached stage 15 (**Figure 3A**). Then explants were treated with DMSO and 2APB until the sibling embryos reached stage 24 (**Figure 3A**). Explants treated with DMSO formed a mucociliary epithelium with MCCs successfully integrated into the superficial epithelial layer as expected (**Figure 3B-C**). In contrast, MCC epithelial integration was defective in explants treated with 2APB (**Figure 3B-C**). The latter indicates that the role of calcium transients in MCC epithelial integration is tissue autonomous.

**Figure 3.**
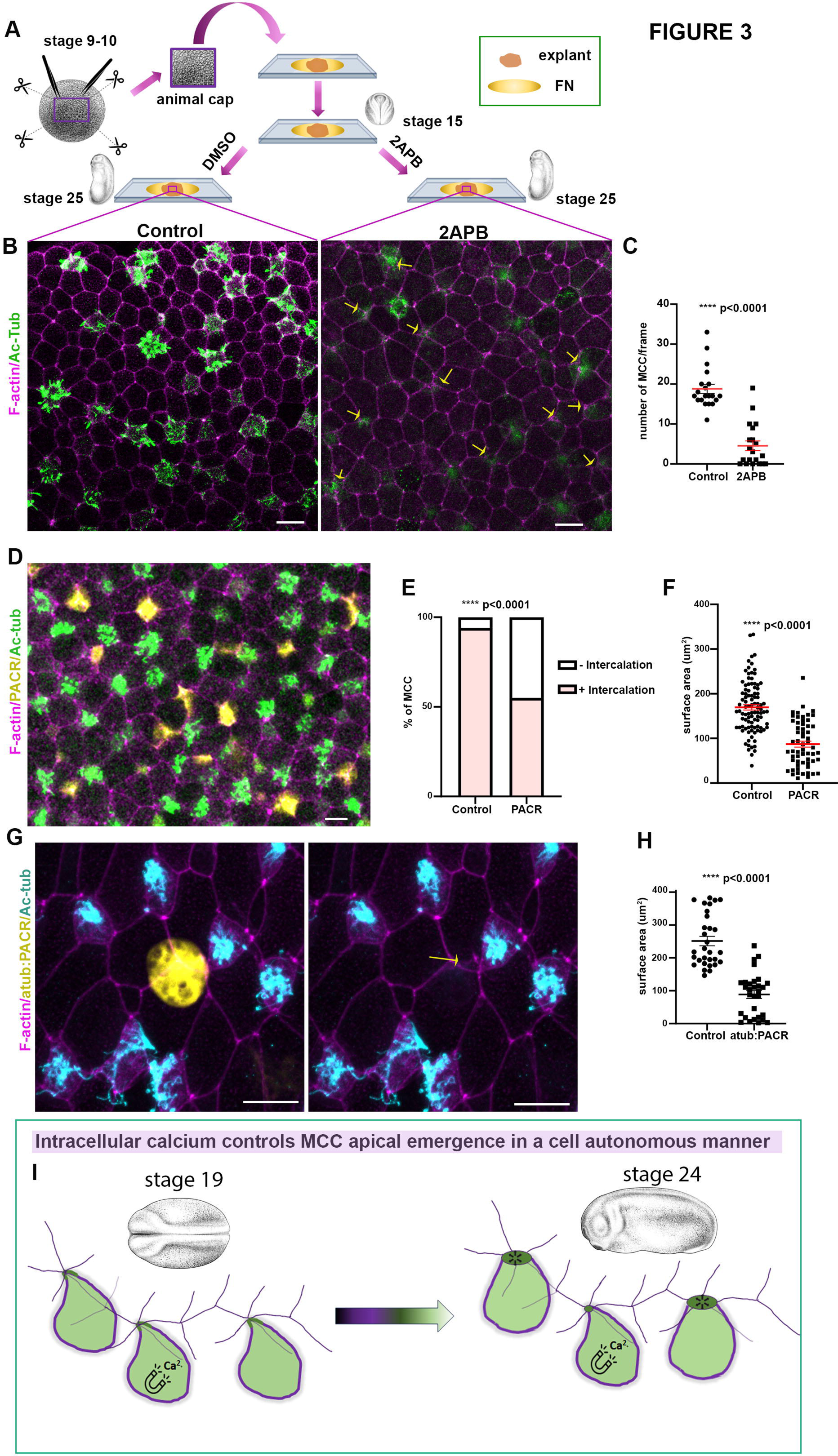
Tissue and cell-autonomous contribution of calcium transients in MCC epithelial integration. A) Diagram depicting the experimental approach followed for the generation of animal-cap explants. B) Representative images of control and 2APB treated animal-caps. Arrows indicate defective epithelial insertion of MCCs in 2APB treated animal-cap. C) Quantification of MCC epithelial integration. n=20 frames from 10 control and 2APB treated explants. Two-sided, unpaired Student’s *t*-test D) Representative image of a stage 24 embryo displaying mosaic expression of PACR- GFP in MCCs. E) Quantification of epithelial integration of control and PACR- expressing multiciliated cells. N=100 control and 108 PACR expressing MCCs, χ^2^ test F) Quantification of the apical cell surface area of control and PACR-expressing MCCs.n=97 control and 59 PACR expressing MCCs. two-sided, unpaired Student’s *t*- test. G) Representative image of skin epithelium from a stage 24 embryos displaying expression of atub:PACR-GFP. MCC expressing atub:PACR-GFP fails to successfully integrate into the superficial epithelium (yellow arrow). H) Quantification of the apical cell surface area of control and atub:PACR-GFP expressing MCCs.n=30 control and 30 atub:PACR expressing MCCs. Two-sided, unpaired Student’s *t*-test I)Schematic showing the cell-autonomous role of intracellular calcium transients in MCC epithelial insertion.

During MCC apical emergence MCCs interact with goblet cells and this interaction is essential for insertion of MCCs in the superficial epithelial layer^5,9,49^. The existence of calcium transients of low amplitude out of our detection limit in goblet cells cannot be excluded. Thus, treatment of 2APB might lead to inhibition of calcium signalling in goblet cells which might be necessary for junction remodelling in these cells, indirectly affecting MCC apical emergence. To exclude the possibility that the defects observed in MCC epithelial integration stem from defects in goblet cells’ junction remodelling we decided to use the genetically encoded Ca^2+^ cage, PACR^50^. PACR is a tool that combines a photosensitive protein domain, LOV2, and a Ca^2+^ binding domain. When PACR is overexpressed in the absence of blue light, Ca^2+^ is bound to PACR causing Ca^2+^ to be sequestered from the cytoplasm^50^. Therefore, in the absence of blue light PACR can be used as a genetically encoded Ca^2+^ chelator. Injection of linear plasmid DNA encoding PACR-GFP led to mosaic expression of PACR in the skin epithelium. Imaging of stage 24 embryos (developed in the dark) in regions where PACR was expressed in MCCs (assessed by increased levels of ac-tub) revealed that PACR+ MCCs displayed defective epithelial integration and apical cell surface expansion (**Figure 3D-F, Movie 5**). To ensure that the defects observed in PACR+ MCCs are not due to ineffective MCC specification, we cloned PACR into a plasmid controlled by the MCC-specific a-tubulin promoter. This allowed us to investigate the effects of calcium chelation specifically after MCC specification and during MCC apical emergence. MCC expressing atub:PACR could not successfully enter the skin epithelium while neighbouring control MCCs successfully entered the epithelium (**Figure 3G-H**). Overall, the above data show that Ca^2+^ is necessary during MCC epithelial integration in a cell-autonomous manner.

### PLC activity is necessary for calcium transient generation and MCC apical emergence

Intracellular calcium transients precede and are necessary for MCC epithelial integration. Inhibition of IP3R-mediated calcium efflux from the ER upon 2APB treatment leads to defective MCC apical emergence (**Figure 4A**). IP3R mediate calcium release is activated by PLC-mediated PIP2 hydrolysis, which leads to the generation of IP3 and DAG (**Figure 4A**)^51,52^. Thus, we hypothesized that PLC activity could regulate calcium mediated MCC epithelial integration. To explore this hypothesis, initially, we sought to monitor PLC activity during MCC epithelial integration. For this we utilize a DAG biosensor, to monitor the dynamics of PLC mediated PIP2 hydrolysis^53^. The biosensor is composed of GFP fused with the C1a domain of PKC-γ that specifically binds to DAG. Live imaging of embryos expressing a GFP-PKC-γ-C1a revealed that PLC displays pulsed activity during MCC apical emergence (**Figure 4B-C, Movie 6**). This supports the hypothesis that PLC is involved in calcium transient generation during MCC epithelial integration. To directly examine the role of PLC for calcium transient generation we first used the PLC-specific inhibitor U73122^54^ and imaged intracellular calcium levels during MCC apical emergence. Live imaging revealed that PLC activity is necessary for calcium transients’ generation in MCCs (**Figure 4D-E**). Next, we went on to examine the role of PLC during MCC epithelial integration. For this we allowed sibling embryos to develop to stage 15 and subsequently embryos were treated with DMSO or U73122 until stage 24. Blockage of PLC activity abrogated MCC apical emergence with MCC failing to enter the superficial epithelial layer or displaying defects in apical cell surface expansion (**Figure 4F-G**). To exclude the possibility that defective MCC epithelial integration stems from the effect of U73122 in other epithelial cell types we decided to use a well- characterized dominant negative construct for PLC, PH-PLCD1^55,56^. Overexpression of PH-PLCD1 blocks PIP2 hydrolysis by PLC through high-affinity binding to PIP2. To assess the role of PLC specifically in MCCs we microinjected DNA of a plasmid construct encoding PH-PLCD1 to achieve mosaic expression. When embryos reached stage 24 we analysed MCC apical emergence in MCC expressing the PH-PLCD1 and surrounded by control epithelial cells. MCC-specific expression of PH-PLCD1 resulted in defective MCC insertion in the skin epithelium mimicking the phenotype induced by the pharmacological inhibition of PLC (**Figure 4H-I**). In summary, our data show that PLC activity is necessary for calcium transient generation in MCCs and is crucial for proper MCC epithelial integration in a cell-autonomous manner.

**Figure 4.**
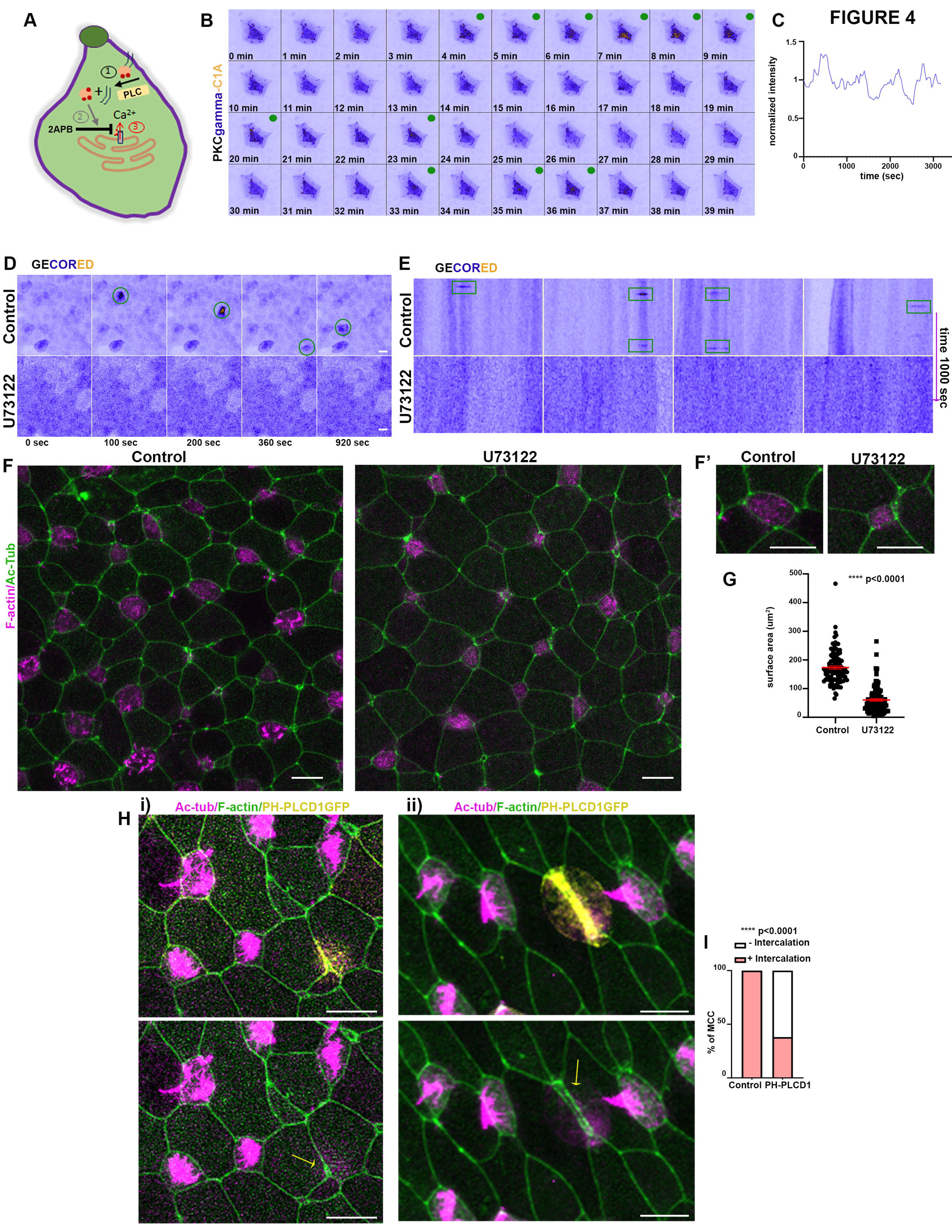
PLC activity is necessary for generation of calcium transients during MCC epithelial insertion. A) Schematic depicting the series of events leading to calcium release from the ER through activation of the IP3 receptor. 1) PLC hydrolyses PIP2 to IP3 and DAG. 2)IP3 binds at the IP3 receptor located at the ER membrane. 3) Activation of the IP3 receptors leads to release of calcium from the ER to the cytoplasm. 2APB inhibits ER mediated calcium release by inhibiting IP3 receptor. B) Stills from a representative time lapse recording showing an intercalating MCC expressing the PLC activity sensor GFP-PKCγ-C1a. PLC displays a transient pulsed activation, as evident by the localized (green dots) at the apical surface of MCC. C)Quantification of GFP-PKCγ-C1 over time. D) Stills from time lapse recordings showing a region of the skin epithelium of control and PLC-inhibitor treated embryos expressing the calcium sensor GECO-RED. Inhibition of PLC suppresses calcium transients (green circles) generation. E) Kymographs from distinct regions of the skin epithelium of control and PLC inhibitor treated embryos. Transient increases in calcium levels in intercalating MCCs (green rectangles) are blocked in presence of the PLC inhibitor. F) Representative images of the skin neuroepithelium of stage 24 control and PLC inhibitor treated embryos. MCCs fail to enter the superficial epithelial (yellow arrows) or expand their surface area (yellow circles) when PLC activity is impaired. F’) Zoomed images of MCCs from a control and PLC inhibitor treated embryo. G) Quantification of MCCs apical surface area in control (n=100 MCCs) and PLC-inhibitor treated embryos (n=100 MCCs). Two-sided, unpaired Student’s *t*-test H) Representative images of the skin epithelium of stage 24 embryos expressing the PLC dominant negative construct PH-PLCD1. Expression of PLCD1 results in defective MCC apical emergence (yellow arrow). I) Quantification of successful MCC apical emergence of control (n=150 MCCs) and PH-PLCD1 (n=116 MCCs) expressing MCCs, χ^2^ test.

### Calmodulin is a downstream effector of Ca^2+^ during MCC epithelial insertion

Our data show that PLC/Ca2+ generated calcium transients regulate Xenopus skin epithelium morphogenesis and specifically MCC apical emergence. Intracellular calcium is a second messenger regulating distinct morphogenetic processes via regulation of various Calcium-binding proteins^27,57^. One well-characterized calcium signalling effector is Calmodulin^58^. Calmodulin(CALM) is composed of four EF-hand domains, each capable of binding a calcium ion. Upon Ca2+ binding, calmodulin undergoes a conformational change which affects its interaction with many proteins^59,60^. Previous studies have shown that calmodulin localizes to flagellar radial spokes^61^. In addition, it has been reported that calmodulin localizes at the centrosomes^62^, a structure similar to basal bodies. To examine the localization of calmodulin in the Xenopus embryo, we used GFP tagged Calmodulin. CALM-GFP displayed centrosomal localization in dividing goblet cells in agreement with previous reports (**Figure S3A**). In MCCs calmodulin displayed axonemal localization, again in agreement with the reported flagellar radial spoke localization (**Figure S3B**)^61^. Additionally, at the apical site of MCCs CALM localized at the basal bodies and the apical actin cytoskeleton (**Figure 5A-C**). CALM basal body localization was evident before MCC apical emergence and apical basal body docking, when basal bodies are in the cytosol (**Figure S3C**). Proteins localized at the basal bodies of MCC have been shown to regulate MCC epithelial integration, thus calmodulin localization is consistent with a role in MCC epithelial integration. To examine if CALM activity is necessary for MCC apical emergence we employed the specific CALM inhibitor, W7. Pharmacological inhibition of CALM resulted in defective MCC apical emergence with MCCs either completely failing to enter the upper epithelial layer or entering the epithelium but failing to expand their apical surface area (**Figure 5D-E**). To examine the role of CALM specifically in MCCs we used a CALM dominant negative construct (CALM1234) that is unable to bind to calcium due to mutations in all four EF-hand domains (D20A, D56A, D93A, and D129A)^63,64^. We injected embryos with plasmid DNA encoding WT-CALM or CALM1234 to achieve mosaic expression. Subsequently, we fixed embryos at stage 2 and analysed MCC expressing WT-CALM or CALM1234 and surrounded by epithelial cells non expressing the exogenous constructs. MCC expressing the WT exogenous successfully integrated in the superficial skin epithelium similarly to control MCCs (**Figure 5F**). On the other hand, MCCs expressing CALM1234 displayed defects in MCC apical emergence (**Figure 5F**). Specifically, a lower percentage of MCC expressing CALM1234 successfully integrated into the epithelial layer, and the cells integrated into the epithelium and expressed CALM1234 had a significantly smaller apical surface area (**Figure 5G-H**). Thus, our data indicate that calcium-bound calmodulin regulates MCC epithelial integration and apical cell surface expansion in a cell-autonomous manner.

**Figure 5.**
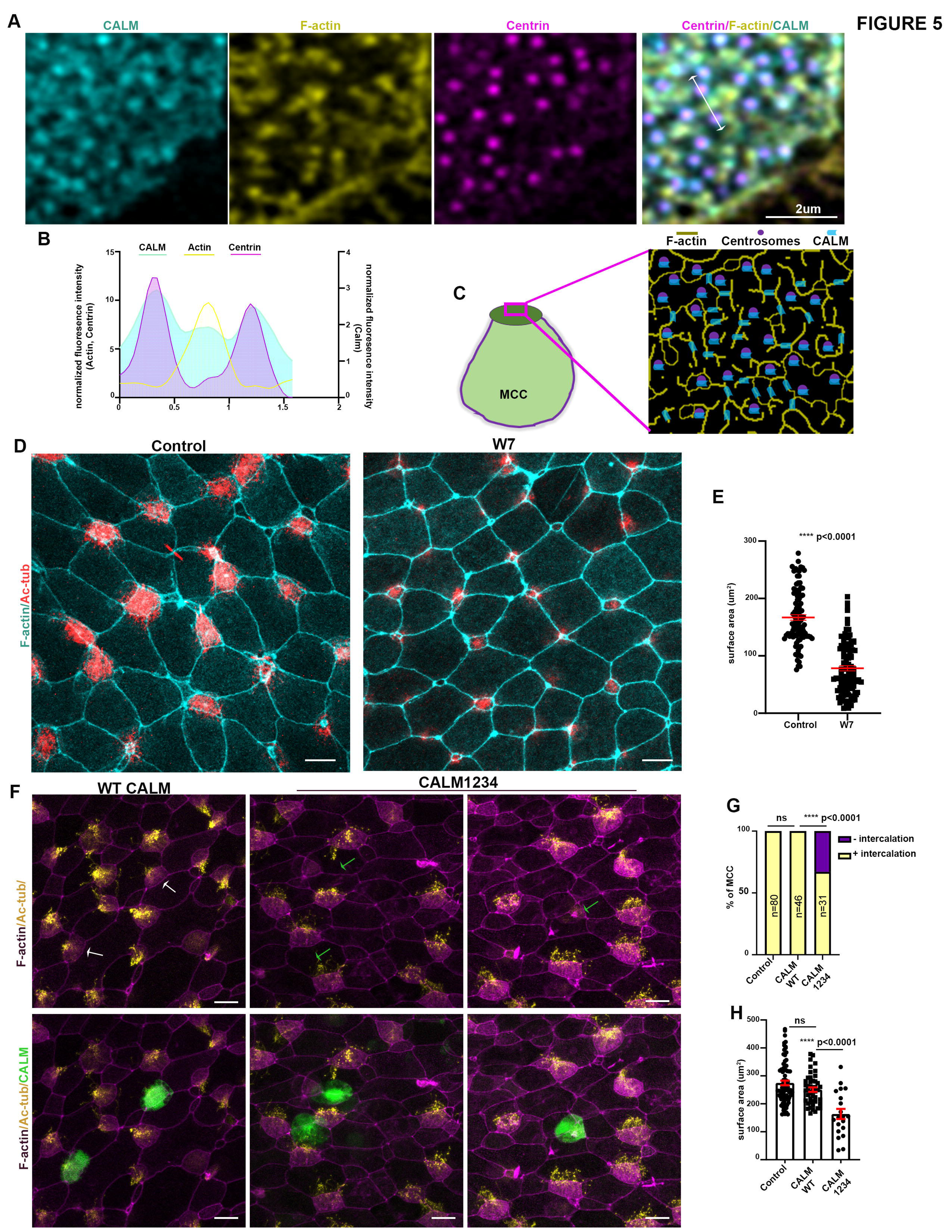
Calmodulin regulates MCC epithelial integration. A) Representative MIP image of the apical surface area of a MCC expressing calmodulin-GFP. Calmodulin displays colocalization with the basal bodies and the apical actin network. B) Fluorescent intensity profile along the double headed arrow in A shows the colocalization of calmodulin with both basal bodies (centrin signal) and the apical actin network. C)Schematic depicting the localization of calmodulin in a MCC in relation to the basal bodies and the apical actin network. D) Representative images of the skin epithelium from control and calmodulin inhibitor treated embryos. MCC fail to fully integrate into the superficial skin epithelium when calmodulin activity is blocked, as evident by their small apical surface area. E) Quantification of the apical surface are of MCCs from control (n=100 MCCs) and calmodulin inhibitor-treated embryos (n=100 MCCs). two-sided, unpaired Student’s *t*-test. F) Representative images of the skin neuroepithelium from embryos expressing wild-type calmodulin or calcium binding deficient calmodulin mutant (CALM1234). Expression of CALM1234 results in defective MCC apical emergence (green arrows) while expression of WT calmodulin does not affect MCC epithelial integration (white arrows). G) Quantification of MCC epithelial integration upon expression of WT and calcium binding deficient mutant calmodulin. χ^2^ test H) Quantification of the apical surface area of MCCs expressing WT and calcium binding deficient mutant calmodulin. N=80 control MCCs, 46 MCCs expressing WT calmodulin and 20 MCCs expressing mutant calmodulin. two-sided, unpaired Student’s *t*-test.

### PLC/Ca^2+^/Calmodulin signalling axis controls apical actin organization in MCC

MCC apical emergence is governed by an apical actin network generating 2D planar pushing forces, essential for epithelial integration and apical cell surface expansion^9,10^. To examine the influence of intracellular calcium in MCC apical network formation during epithelial integration we performed live imaging of tailbud embryos expressing the calcium sensor GECO-RED with Utr-GFP allowing visualization of F-actin. Live imaging revealed that an increase in intracellular calcium precedes the enrichment of apical actin (**Figure 6A-C, Movie 7**). This suggests a possible functional role of calcium transients in the establishment of the apical F-actin network in these cells. To examine if intracellular calcium increase is sufficient to induce apical actin enrichment, we decided to pharmacologically elevate intracellular calcium using thapsigargin (THA)^27^. Live imaging of embryos expressing atub:GECORED/atub:UtrGFP immediately after the addition of THA to the culture medium revealed that an increase of intracellular calcium led to the enrichment of the F-actin at the apical region of MCCs (**Figure 6D- E, Movie 8**). This further suggests that intracellular calcium and its downstream effectors might be necessary for the correct formation of the apical actin network during MCC intercalation. Treatment of embryos with 25um 2APB completely blocks MCC epithelial integration, with MCC failing to acquire an apical surface (**Figure 2A-B**). Thus, to explore the role of calcium transients on apical actin network formation, we treated embryos from stage 15 onwards with 12.5uM 2APB. Embryos were fixed and analysed at stage 24. Treatment of embryos with 12.5uM of 2APB did not completely block MCC epithelial integration but significantly affected MCC apical cell surface area expansion (**Figure 6F-G**). In addition, the apical actin network formation was defective in MCC of embryos treated with 2APB, as evident by the reduction in apical actin intensity (**Figure 6F,H**). In agreement with the above, apical actin network establishment was also impaired in MCCs expressing the genetically encoded calcium chelator PACR (**Figure 6I-K**). Subsequently, to examine the role of PLC and CALM in apical actin network establishment, we used the inhibitors U73122 and W7 respectively. Specifically, embryos were treated from stage 15 onwards with either U73122 or W7 and allowed to develop to stage 24. Immunofluorescence analysis revealed that PLC and CALM activity is necessary for the proper establishment of the apical actomyosin network during MCC apical emergence (**Figure 6L-O**). Overall, these data strongly indicate that the PLC/Ca^2+/^Calm signalling axis regulates MCC epithelial integration by regulating the establishment of the apical actin network.

**Figure 6.**
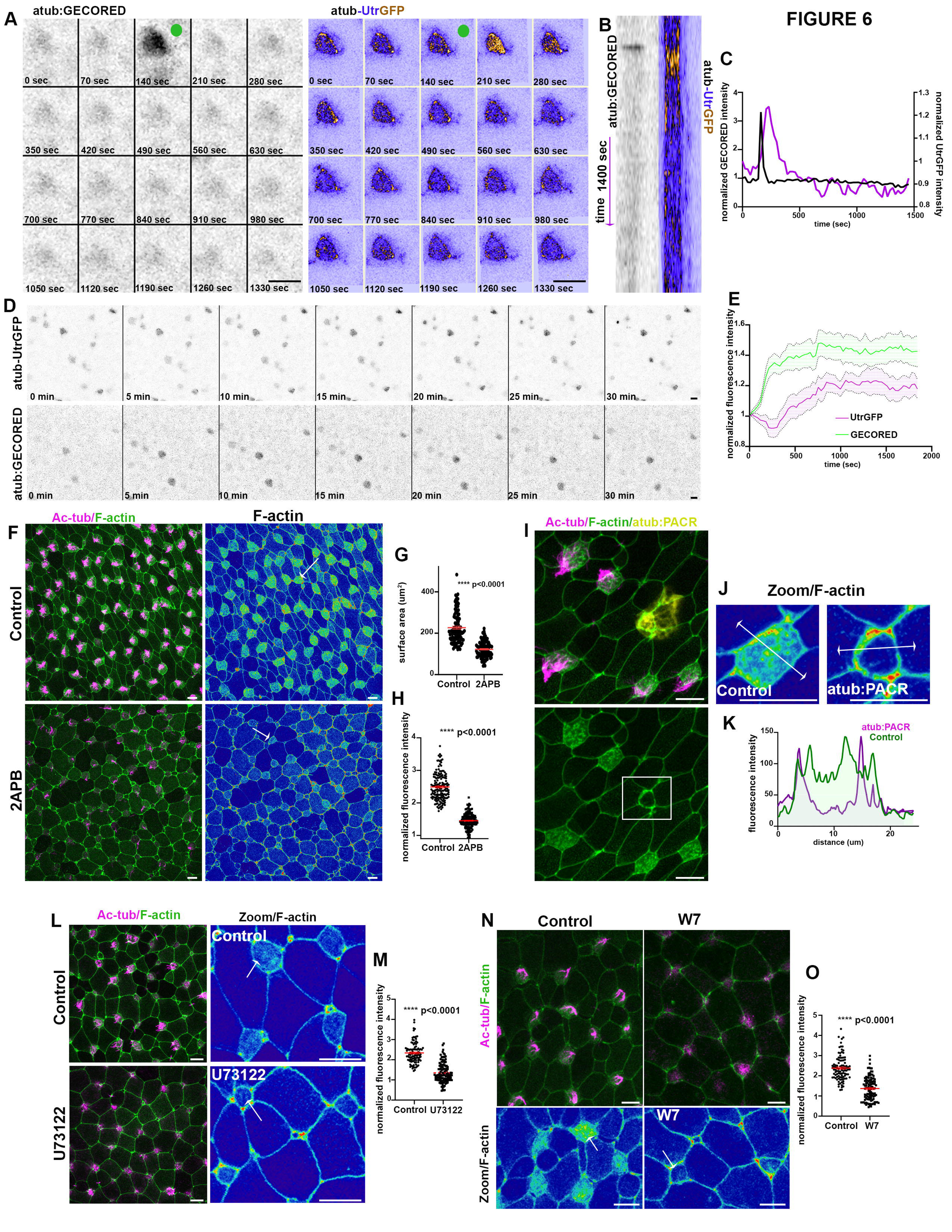
PLC/Ca^2+^/Calmodulin signalling governs apical actin cytoskeleton organization in MCC. A) Stills from a time-lapse recording of an intercalating multiciliated cell expressing atub:GECORED and atub:UtrGFP. Calcium transient (green dot) is followed by an enrichment of the apical actin. B) Kymograph for the MCC shown in A showing that a calcium transient precedes a temporary enrichment of the apical actin network. C) Quantification of GECO-RED and Utr-GFP fluorescent intensity over time. D) Stills from a time lapse recording of an embryo expressing atub: GECORED and atub:UtrGFP and treated with thapsigargin. Thapsigargin treatment led to elevation of calcium levels in MCCs with a concomitant enrichment of the apical actin network. E) Quantification of GECORED and UtrGFP fluorescent intensity over time for 10 MCCs from D. Elevation of intracellular calcium in MCCs leads to enrichment of the apical actin network, F) Representative MIP images of the skin epithelium from control and 2APB treated embryos. Fluorescent intensity coded imaged show defective formation of the apical actin network in 2APB treated embryos (white arrows). G) Quantification of apical surface area of MCCs in control embryos and embryos treated with 12.5uM 2APB. N=100 MCCs; two-sided, unpaired Student’s *t*-test. H) Quantification of actin apical enrichment in MCCs from control and 12.5uM 2APB treated embryos. n=160 MCCs; two-sided, unpaired Student’s *t*-test. I) Representative image showing defective apical actin enrichment in a MCC expressing PACR-GFP (white square). J) Fluorescent intensity color coded images of a control and a PACR expressing MCC. K) Fluorescent intensity profile along the double headed arrow in I reveals the defective formation of the apical actin network in PACR expressing MCC. L) Representative image of the skin epithelium of control and PLC inhibitor treated embryos. Fluorescent intensity color coded images show defective formation of the apical actin network in the absence of PLC activity (white arrows). M) Quantification of actin apical enrichment in MCCs from control and PLC inhibitor treated embryos. n=100 MCCs from control embryos and 120 MCCs from PLC- inhibitor treated embryos; two-sided, unpaired Student’s *t*-test. N) Representative image of the skin epithelium of control and calmodulin inhibitor treated embryos. Fluorescent intensity color coded images show defective formation of the apical actin network in the absence of calmodulin activity (white arrows). O) Quantification of actin apical enrichment in MCCs from control and calmodulin inhibitor treated embryos. =100 MCCs from control embryos and 110 MCCs from calmodulin inhibitor treated embryos; two-sided, unpaired Student’s *t*-test

### Intracellular calcium regulates MCC apical actin network stability

Apical actin network establishment and MCC epithelial integration are regulated via multiple pathways and processes. Specifically previous work demonstrated that apicobasal polarity establishment^7,21^, RhoA activity^10^, basal body number^22^ and actin network polymerization^9^ and stabilization^24^ are all involved in MCC apical emergence. Therefore, we went on to explore the input of calcium signalling in the above processes during MCC epithelial integration. To examine the effect of calcium signalling on apicobasal polarization we injected embryos with plasmid DNA encoding PAR3-GFP and subsequently assessed PAR3 localization in MCC of control and 2APB treated embryos. Par3 apical enrichment was unaffected by 2APB treatment (**Figure S4A**). This indicates that MCC apicobasal polarity is not affected by the disruption of calcium transients. To further asses the establishment of apicobasal axis in the absence of calcium transients we assessed basal body localisation prior to MCC epithelial insertion. Basal bodies were apically localized in MCC of control and 2APB treated sibling embryos (**Figure S4B**). These data show that MCC apicobasal polarization does not require calcium signalling.

RhoA activity is necessary for the generation of 2D pushing forces during MCC apical cell surface expansion^10^. To examine if calcium transients affect RhoA activity we used the RhoA activity sensor rGBD^65^. During MCC intercalation active RhoA localizes at the basal bodies^65^. Active RhoA localization and basal body docking were not affected in embryos treated with 12.5um of 2APB, even though these cells displayed defects in apical cell surface area expansion (**Figure 7A**, **Figure 6G**). Thus, Rhoa activation during MCC apical emergence is calcium independent.

**Figure 7.**
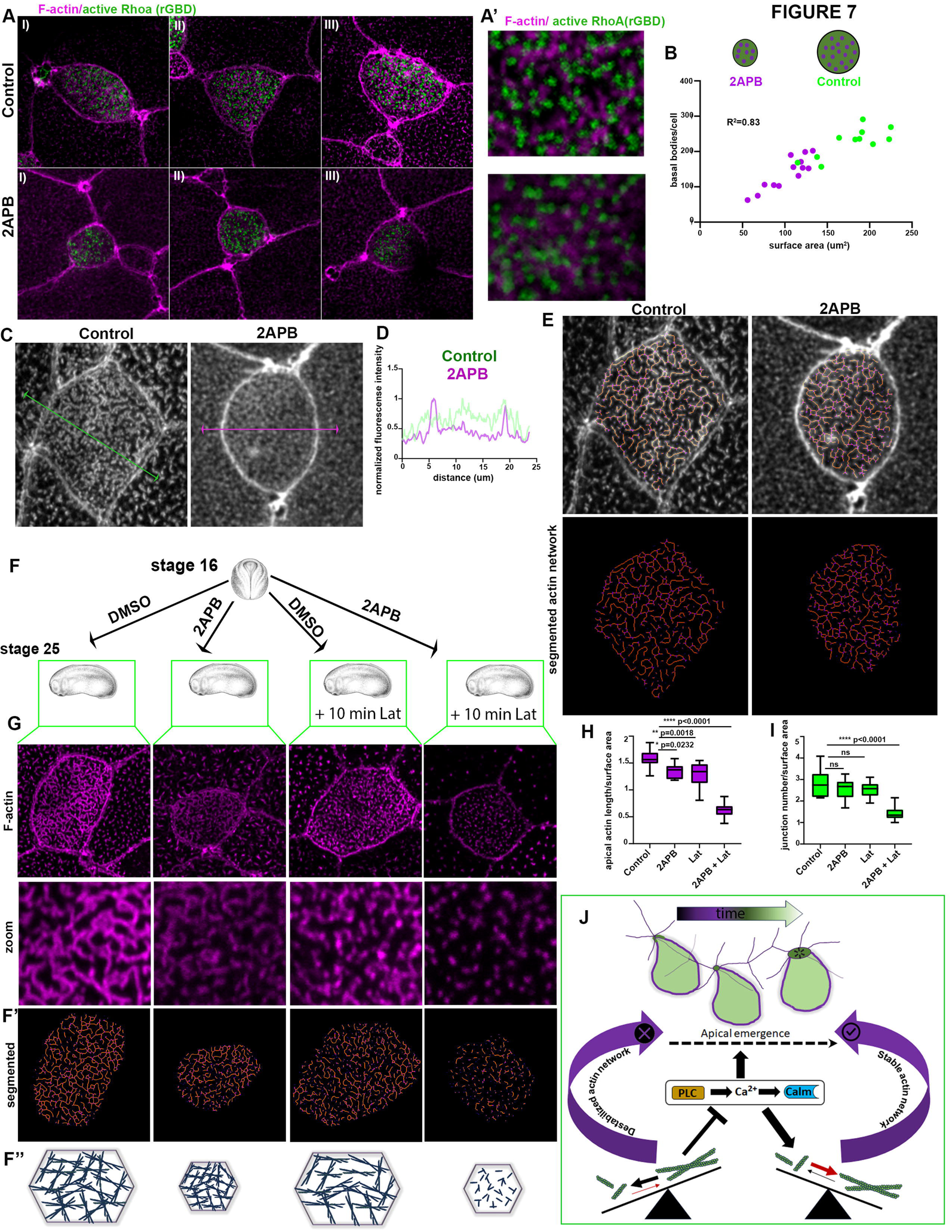
Intracellular calcium controls the stability of the apical actin cytoskeleton in MCC. A) Representative images showing active-RhoA localization in MCCs of control and 2APB treated embryos. A’) Zoomed images showing that the association of active-RhoA with basal bodies is not affected in embryos treated with 2APB. B) Correlation plot between basal body number and apical surface area of MCCs in control and 2APB treated embryos. While MCCs cells display a lower number of MCCs, basal body number is positively correlated with the apical surface are of MCCs. C) Representative MIP images of the apical actin network of MCCs from control and 2APB treated embryos. D) Fluorescent intensity profile along the double headed arrows in C show that 2APB treatment results in defective enrichment of the apical actin network. E) Segmentation of the of the apical actin network of MCCs reveals that 2APB treatment does not affect the architecture of the apical actin network. Orange: actin filaments. Purple: Actin filament junction points. F) Schematic depicting the experimental design to assess the impact of 2APB treatment on MCC apical actin network stability. G) Representative MIP images of the apical actin network in control, 2APB treated, Lat treated and 2APB+Lat treated MCCs. G’) Segmented images of the apical actin cytoskeleton showing that 10min Lat exposure results in the disruption of the apical actin cytoskeleton of MCCs only when embryos are exposed to 12.5uM 2APB. G’’) Schematics summarizing the status of MCC apical actin network in different conditions. H) Quantification of the total apical actin network normalized against the apical surface area. While treatment of embryos with 12.5uM 2APB and 10min Lat has a minimal effect on the total actin network length, combination of these treatments results in destruction of the apical actin network. I) Quantification of total junction points of the apical actin network normalized against the surface area showing that the networks architecture is only lost only when 12.5um 2APB and 10min Lat treatments are combined. I) Proposed model for the action of PLC/Ca^2+^/Calmodulin signalling axis during MCCs epithelial integration.

Abrogation of centriole amplification and a lower number of basal bodies results in defective MCC apical emergence^22^. Thus, we went on to examine the effect of intracellular calcium on centriole amplification. For this we analysed the number of basal bodies in sibling control embryos and embryos treated with 12.5 2APB. The total number of basal bodies was lower in 2APB-treated embryos (**Figure 7B**). However, MCCs in these embryos have a smaller apical surface area (**Figure 6G**, **Figure 7A**). It has been reported that the number of basal bodies is scaled to the apical surface area through a mechanosensitive mechanism and a minimum apical surface area can be achieved independent of centriole amplification^11^. Importantly, the number of basal bodies displayed a high correlation with the size of the apical cell surface area both in control and 2APB-treated embryos (**Figure 7B**). Thus, our data suggest that the defects in MCCs epithelial integration upon inhibition of calcium transients do not stem defective centriole amplification.

Since apicobasal polarity establishment, RhoA activation and basal bodies amplification are not affected by the inhibition of calcium transients we went on to examine how intracellular calcium affects the apical actin network during MCC epithelial integration. For this initially we acquired high-resolution images of the apical actin cytoskeleton in control MCCs and MCCs treated with a suboptimal amount of 2APB (12.5uM). The enrichment of the apical actin network was defective in MCCs of embryos treated with 2APB, as described above (**Figure 7C-D**). In contrast, the overall architecture of the network (overall length and junctions) did not display major defects (**Figure 7E**). This led us to hypothesize that the initial establishment of the actin network can take place in the absence of calcium transients, but its maintenance depends on calcium transient generation. The F-actin network is a dynamic structure that is constantly assembled and disassembled from G-actin monomers. Since apical actin enrichment is severely affected upon abrogation of PLC/Ca^2+^/CALM signalling, while the F-actin network architecture is not affected, we postulated that this signalling axis restricts the pace of actin disassembly. To explore this possibility, we employed Latrunculin A (LatA), which binds to G-actin in a stoichiometric 1:1 ratio and limits actin polymerization^66^. The rate of actin network disassembly can be assessed by subjecting MCCs to low concentration of LatA for a predetermined period and monitoring the effects on the apical actin network. We administered a suboptimal concentration of 2APB (12.5uM) for this experiment to ensure apical emergence of MCCs. Next, we exposed the control and 2APB treated sibling embryos to 2uM LatA or DMSO for 10 minutes (**Figure 7F**). The apical actin network in MCCs was examined via segmentation of the apical actin network. Both the normalized apical actin network length and normalized number of junctions in the actin network were severely affected only in embryos treated with 2APB+ Lat (**Figure 7H-I**). This experiment shows that the apical actin network stability is drastically reduced upon inhibition of calcium signalling (**Figure 7G**). Thus, MCC epithelial integration defects upon abrogation of normal intracellular calcium homeostasis stem from destabilization of the apical actin network.

## Discussion

Insertion of basally located progenitors into an existing epithelial layer is essential for tissue morphogenesis and homeostasis. Basal stem cell integration into an epithelial layer necessary for tissue homeostasis has been documented in the mammalian airway epithelium^4^, olfactory epithelium^15^, cornea^18^, prostate^16^ and the Drosophila midgut^17,19^. Additionally, this morphogenetic behaviour has been reported during embryogenesis. Single-cell or multicellular epithelial integration contributes to endoderm and node formation in mammals^12–14^, and mucociliary skin epithelium development in Xenopus^5,6^.

In this work, we focused on the role of intracellular calcium transients in epithelial integration of MCCs during Xenopus mucociliary skin epithelium development. Our data show that intracellular calcium transients precede MCC apical emergence and are necessary for MCC epithelial integration in a tissue and cell- autonomous manner. These calcium transients are generated through PLC activity, with calmodulin acting as an effector of intracellular calcium during MCCs cell integration. Mechanistically, our work reveals that the PLC/Ca^2+^/Calmodulin signalling axis regulates MCC epithelial insertion by contributing to apical actin cytoskeleton network stability. We propose that defects in the PLC/Ca^2+^/Calmodulin signalling pathway result in inadequate stabilization of the apical actin network, causing its disruption and impairing the apical actin-generated pushing forces essential for MCCs epithelial integration (**Figure 7J**).

Our work reveals that intracellular calcium transients precede MCC epithelial integration. Using surface ectoderm explants and a genetically encoded calcium chelator we further showed that calcium transients regulate MCC apical emergence in a tissue and cell-autonomous manner. In addition, our data show that not all steps of MCC epithelial integration are Ca^2+-^dependent. The initial intercalation of basally located MCC progenitors through the basolateral domain of apical-located epithelial goblet cells^5,6^ and the mechanical probing of apical junctions by MCCs^8^ are calcium- independent. These results indicate that calcium transients are not essential for basolateral junction remodelling and filopodia generation necessary for initial intercalation of MCCs^44^. In contrast, our data show that calcium is required for epithelial integration and MCC apical surface area expansion^9^.

The processes regulated by intracellular calcium during MCC epithelial integration are controlled by RhoA-mediated polymerization of the apical actin network^10,28^. Specifically, polymerization of the apical actin network has been shown to generate 2D pushing forces at the apical site of MCCs^9^, necessary for epithelial integration and subsequent expansion of the surface area. Our data show that the PLC/Ca2/Calmodulin signalling axis is necessary for apical actin network enrichment. This is consistent with the fact that only epithelial integration and MCC apical surface area expansion are calcium dependent.

Calcium transients during MCC epithelial integration require PLC activity for their generation. This suggests that PIP2 concentration in MCCs will be important in this process, since PIP2 hydrolysis by PLC generates IP3 necessary for calcium release from ER stores^51,52^. In addition to IP3, PIP2 hydrolysis results in the generation of DAG. In this work, we don’t study the input of DAG in MCC apical emergence. However, it is possible that DAG production could regulate MCC epithelial integration or other processes during mucociliary epithelium development through the activation of conventional or novel PKC^67^.

Intracellular calcium is necessary for efficient apical actin network stabilization during MCC epithelial integration. Ca^2+/^Calmodulin regulate actin network architecture and dynamics via interaction with several actin binding proteins^30^. Here we show that calmodulin becomes associated with the basal bodies and the apical actin network in MCCs. This localization of Calmodulin is consistent with a role in the regulation of actin network stability. Calcium-bound calmodulin has been shown to interact with actin- regulating proteins such as INF2^33^, IQGAP1^32^, and COBL^31^. COBL has been shown to display enriched expression in ependymal MCCs regulating the apical actin network^68^, in agreement with the role of Ca^2+/^Calmodulin signalling axis in Xenopus MCCs.

In conclusion, our findings highlight the critical role of intracellular calcium transients during epithelial integration of MCCs during Xenopus mucociliary skin epithelium development. Epithelial integration of basally located progenitors has been documented during tissue morphogenesis and homeostasis contributing embryogenesis and adult organ function. Thus, understanding the molecular mechanisms behind calcium-mediated MCC integration provides valuable insights into broader processes of epithelial morphogenesis and tissue homeostasis.

## Materials and Methods

### Xenopus embryos, microinjections and cloning

Female adult *Xenopus laevis* frogs were induced to ovulate by injection of human chorionic gonadotropin. Eggs were fertilised *in vitro*, after acquisition of testes from male frogs, dejellied in 2% cysteine (*pH* 7.8) and subsequently reared in 0.1× Marc’s Modified Ringers (MMR). mRNA and microinjection. For microinjections, embryos were placed in a solution of 4% <u>Ficoll</u> in 0.33× MMR and injected using a glass capillary pulled needle, forceps, a Singer Instruments MK1 micromanipulator and Harvard Apparatus pressure injector at the 4-cell stage. After injections, embryos were reared for 1 hour in 4% Ficoll in 0.33× MMR and then washed and maintained in 0.1× MMR. Injected embryos were allowed to develop early tailbud stage (Nieuwkoop and Faber stage 19) at 17°C and imaged live or allowed to develop to the appropriate stage and then fixed in 1x MEMFA for 2 hours at room temperature. Capped mRNAs encoding were *in vitro* transcribed using mMessage machine kits (Ambion). The amount of mRNA per 4nl of microinjection volume was as follows: membrane-GFP, 100 pg; GECO-RED, 150pg; rGBD, 100pg; RFP-Centrin:80pg. For microinjection of DNA constructs, atub:UtrGFP, atub:GECO-RED, PACR (Addgene #55774), atub:PACR, CALM-GFP (Addgene #47602), CALMWT (Addgene #111499), CALM1234 (Addgene #111518). Par3GFP, GFP-PKC-γ-C1a(Addgene #21205), PH-PLCD1 (Addgene #21179) we injected 80pg of DNA per blastomere. The atub:GECORED and atub:UtrGFP constructs were generated by replacing the UtrophinGFP coding sequence in atub:UtrGFP plasmid with the coding sequences of GECORED and GFP- PACR using the In-fusion cloning kit.

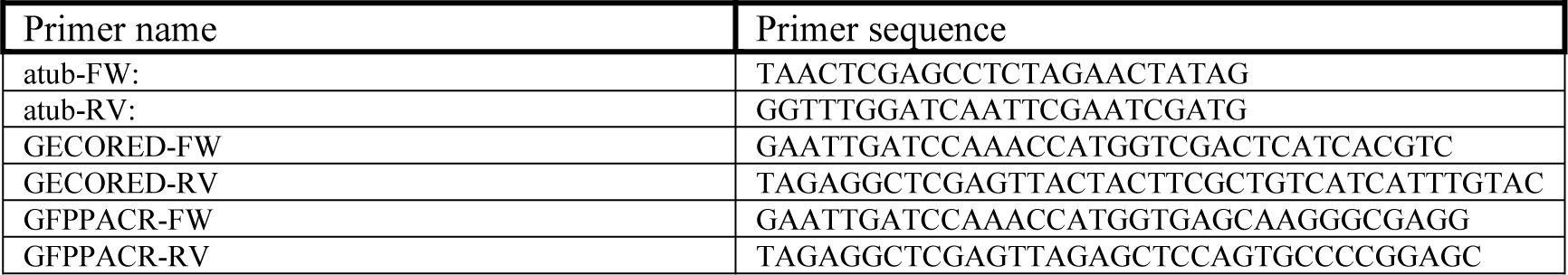

### Immunofluorescence

Immunofluorescence was performed as previously described^57^. Briefly, embryos were fixed for 2 hours at room temperature with MEMFA, permeabilized in PBST (1 × PBS, 0.5% Triton, 1% dimethyl sulfoxide) and blocked for 1 hour in 10% donkey serum. For ZO-1 staining embryos were fixed in methanol over-night at -20 and before blocking the embryos were rehydrated with serial dilution of methanol/PBS. Primary antibodies were incubated overnight at 4 °C. We used a primary antibody against acetylated tubulin, β-catenin (1:500, 11279-R021, SinoBiological), E-cadherin (1:100,5D3, DSHB) and ZO-1 (1:200, 21773-1-AP, Proteintech). Embryos were washed in PBST and incubated for 2 h with secondary antibodies at RT, washed several times and post-fixed in 1XMEMFA. Secondary antibodies used were Alexa fluor 488 (1:500, Invitrogen), Phalloidin was incubated together with the secondary antibodies, and phalloidin 647+(1:500, Invitrogen).

### Ectoderm explants

Animal caps were dissected at stage nine as described previously (Werner and Mitchell, 2013). Subsequently the animal caps were transferred on a slide treated with fibronectin (25 μg/ml) and culture in Danilchik’s for *Amy* (DFA) media supplemented with antibiotic/antimycotic until sibling embryos reached to stage 25.

### Inhibitor Treatment

Embryos were treated from stage 15 to stage 24 with 2APB (25uM and 12,5uM), U73122 (2.5uM) and W7 (50uM). Before treatment with the inhibitors the vitelline membrane was mechanically removed, and embryos were allowed to recover for at least 1 hour. For Lat treatment stage 24 embryos were treated with 10uM Lat for 10 minutes when embryos were fixed.

### Live imaging

Live imaging of Xenopus embryos was performed on a ZEISS LSM 710 confocal microscope with a Plan-Apo 40X, NA 1.1 or a Plan-Apo 25X, NA 0.8 objectives. The ZEISS ZEN software was used during imaging. In order to prevent the coverslip from pressing against the epidermis, embryos were imaged in a customized chamber composed of a thick layer of vacuum grease on a microscope slide. During imaging, embryos were stored at room temperature and mounted in 0.1× MMR.

### Image analysis

All image analysis and quantification were carried out using Fiji software^69^. To analyze apical actin network structure, we followed an automated segmentation protocol previously described^70^. The total length of the apical actin network and the number of junctions in the actin network was normalized by dividing the values of total network length and total junction with the surface area of each MCC. For quantification of MCC intercalation we quantified the number of successfully intercalated MCCs in images with the same frame size (212.55 x212.55 um).

### Statistics

GraphPad Prism 8.0 software was used for all statistical analysis performed. The sample size of the experiments carried out was defined based on previous experimental experience. Quantitative data presented, show the mean ± s.e.m., or the total number of datapoints obtained. The statistical tests carried out on the quantitative data obtained are annotated in each figure legend.

## Author contributions

Conceptualization: N.C.; Methodology: N.C.; Investigation: N.C.; Writing - original draft: N.C.; Writing - review & editing: N.C., P.A.S.; Funding acquisition: N.C., P.A.S.

### Acknowledgements

We thank Dr Peter Walentek for kindly providing the atub:UtrGFP plasmid.

