## Supplemental Figures for "Calcium transients regulate epithelial integration of multiciliated cells during Xenopus skin development"

### Supplementary Figure legends:

**Supplementary Figure 1**

A) Kymographs of single MCCs expressing GECO RED. Each multiciliated cells displays at least one calcium transient. B) Quantification apical surface area of single MCCs. Red bars represent calcium transients. Increase of the apical surface area is preceded by calcium transients. C) Stills from a time-lapse recording of a stage 19 embryo expressing atub:GECORED and treated with 2APB. 2APB treatment results in abolishment of calcium transients in MCCs. D) Kymographs of representative MCCs from D showing the blockage of transient calcium level increases in embryos treated with 2APB. E) Quantification of calcium transient events in MCCs from control and 2APB treated embryos. $\chi^2$ test $p<0.0001$.

**Supplementary Figure 2**

A) Representative images of the skin epithelium of control and 2APB treated embryos stained with β-catenin. B) Fluorescent intensity profile along the double headed arrows in A showing the enrichment of β-catenin at cell-cell junctions. C) Representative images of the skin epithelium of control and 2APB treated embryos stained with E-cadherin D) Fluorescent intensity profile along the double headed arrows in A showing the enrichment of E-cadherin at cell-cell junctions. C) Representative images of the skin epithelium of control and 2APB treated embryos stained with ZO-1. D) Fluorescent intensity profile along the double headed arrows in A showing the enrichment of ZO-1 at tight junctions.

**Supplementary Figure 3**

A) Localisation of GFP-Calmoduin in a diving skin epithelial cell. Magnified image revealed strong centrosomal localization. B) Localization of GFP-calmodulin in MCCs of stage 32 embryo. Calmodulin displays cilia axonemal localization. B) Localization of GFP-calmodulin in MCCs of stage 18 embryo. Calmodulin displays basal body localization before the anchoring of basal bodies at the apical cell surface.

**Supplementary Figure 4**

A) Representative images of stage 24 control and 2APB treated embryos expressing Par3-GFP. Fluorescent intensity profiles along the apicobasal axis of MCCs reveal that Par3 apical localization is unaffected by 2APB treatment. B) Representative images of MCCs from stage 23 control and 2APB treated embryos expressing the basal bodies marker centrin. XZ and YZ projections show that even though MCCs in 2APB-treated embryos do not acquire apical surface, basal bodies in these cells are localized apically.

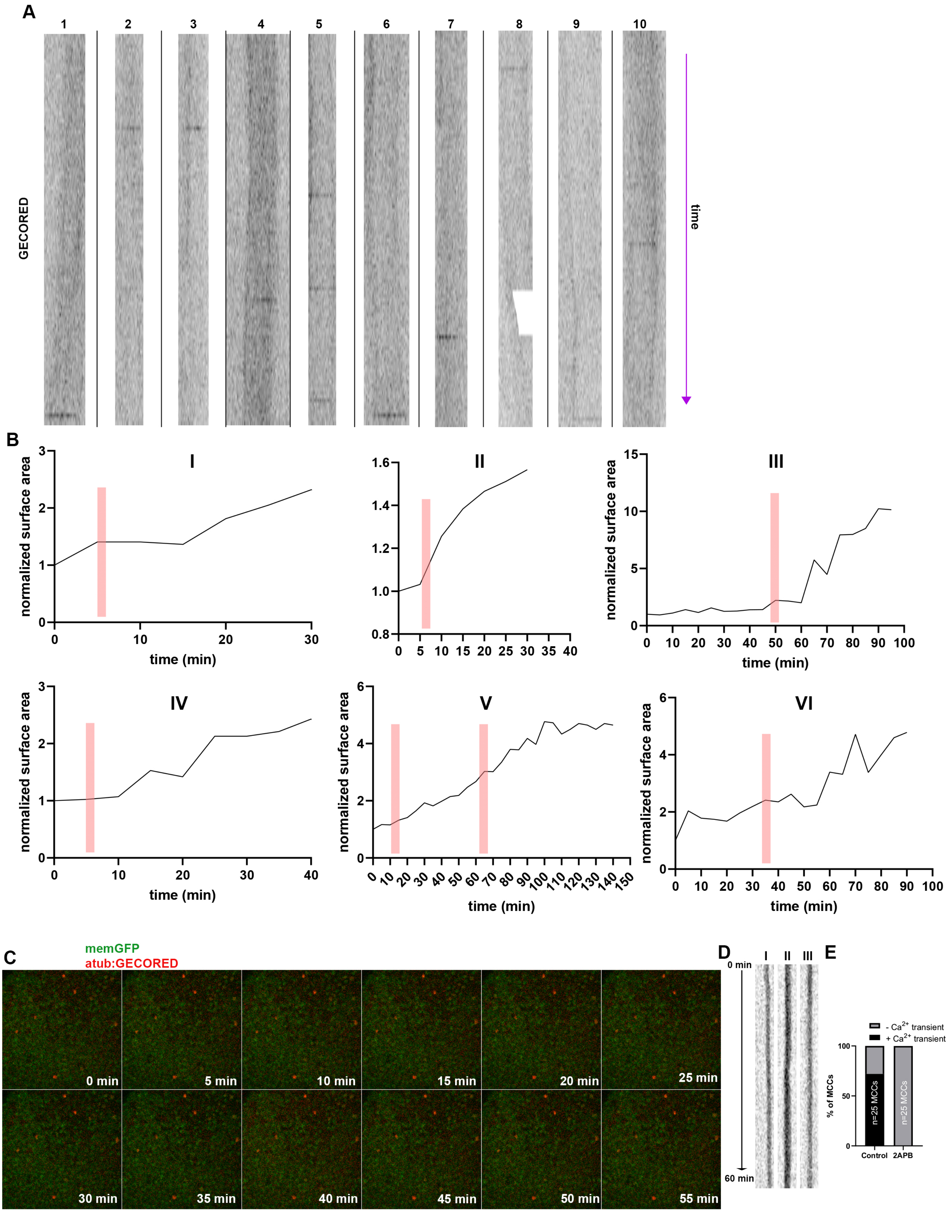

A
1
2
3
4
5
6
7
8
9
10
GECORED
time
B
I
II
III
IV
V
VI
normalized surface area
time (min)
C
memGFP
atub:GECORED
0 min
5 min
10 min
15 min
20 min
25 min
30 min
35 min
40 min
45 min
50 min
55 min
D
0 min
60 min
E
- Ca2+ transient
+ Ca2+ transient
% of MCCs
n=25 MCCs
n=25 MCCs
Control
2APB

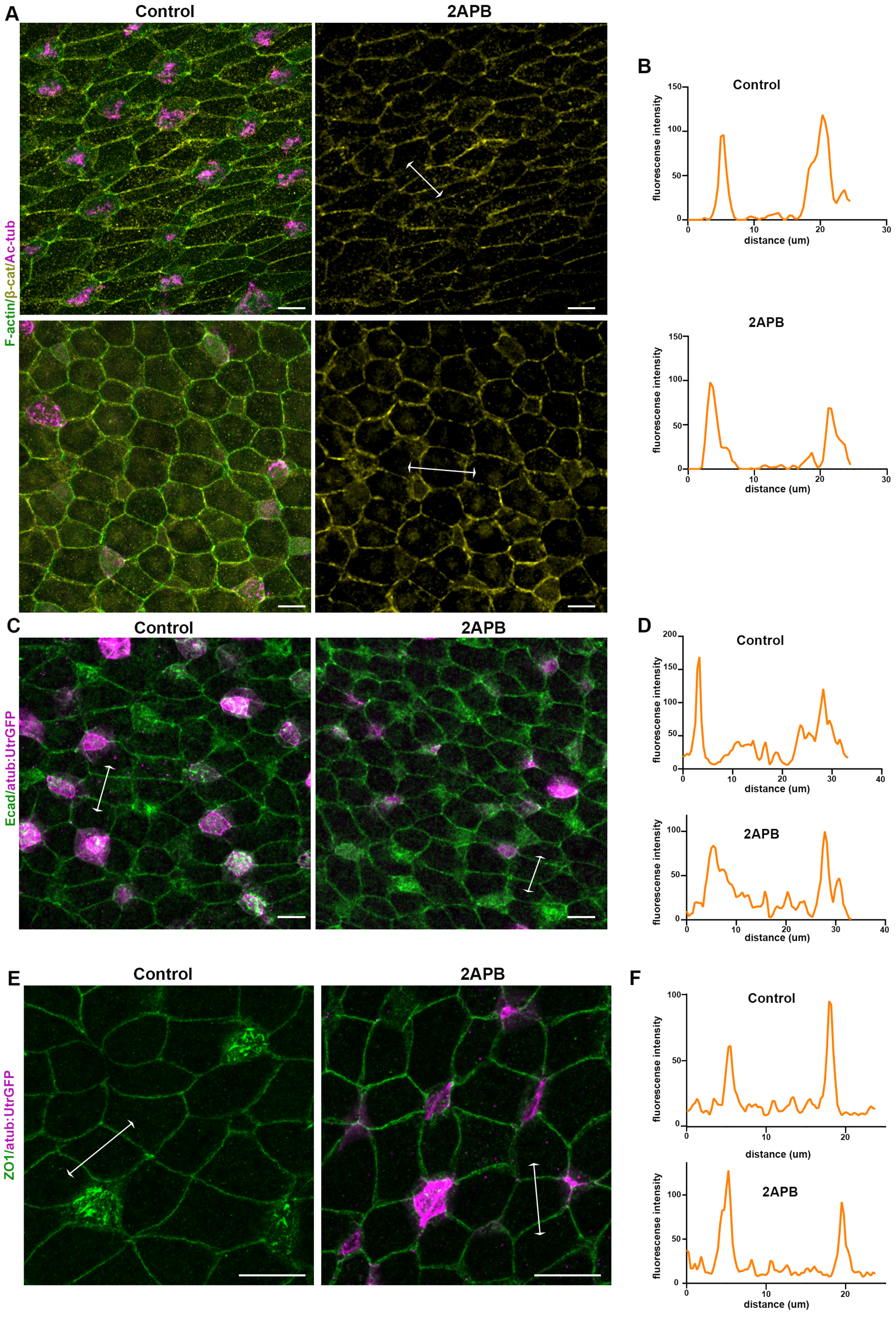

A
Control
2APB
F-actin/β-cat/Ac-tub
B
Control
fluorescense intensity
distance (um)
2APB
C
Control
2APB
Ecad/atub:UtrGFP
D
Control
2APB
E
Control
2APB
ZO1/atub:UtrGFP
F
Control
2APB

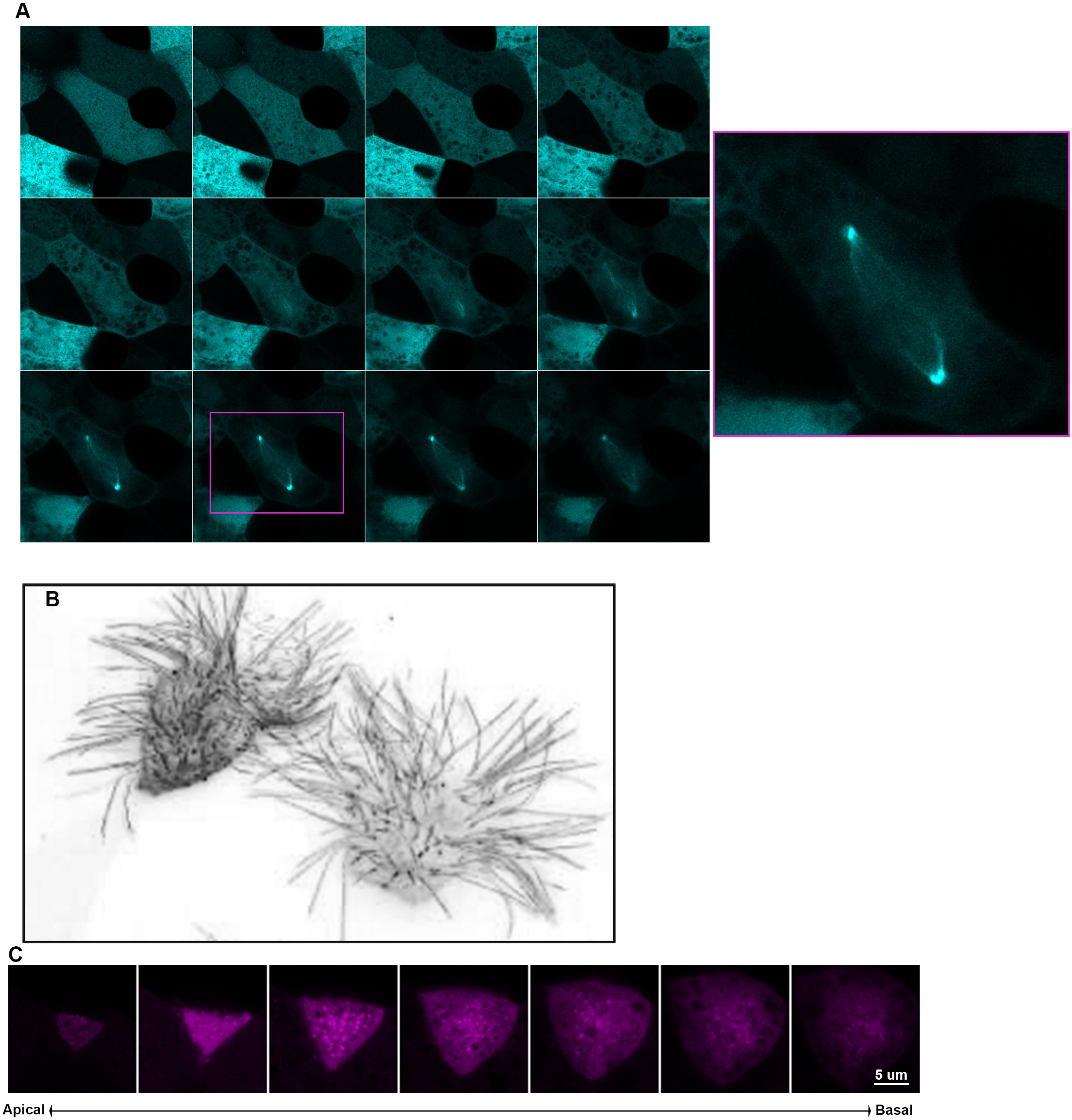
A
B
C
5 um
Apical
Basal

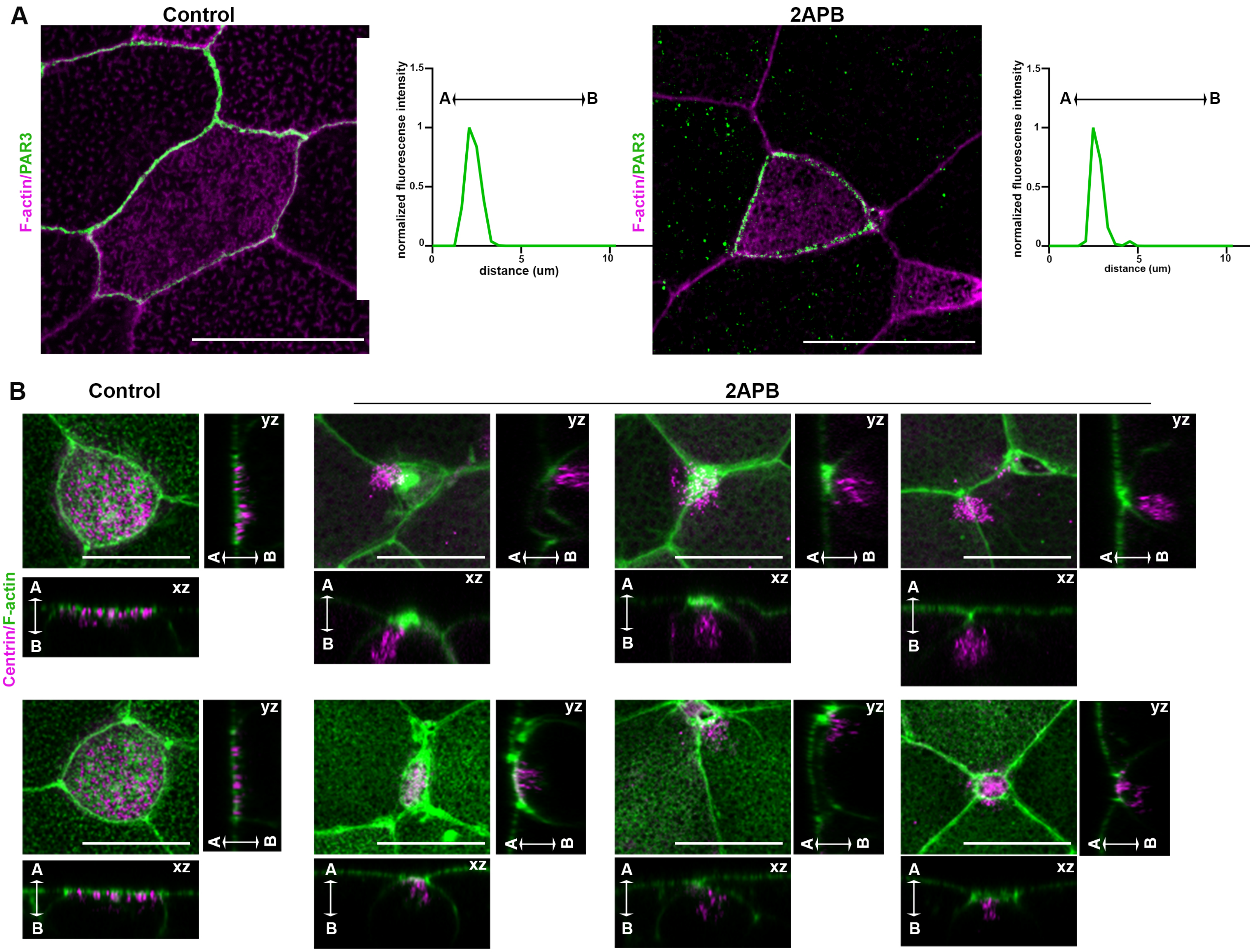
A
Control
2APB
F-actin/PAR3
normalized fluorescense intensity
distance (um)
A
B
1.5
1
0.5
0
5
10
B
Control
2APB
Centrin/F-actin
yz
xz
A
B
